# SOX10 Expression Levels may be a Critical Mediator of White Matter Dysfunction in Schizophrenia

**DOI:** 10.1101/2022.06.14.496174

**Authors:** Donna L McPhie, Boyu Ren, Juan Antonio Garcıa-Leon, Catherine M Verfaillie, Bruce M Cohen

## Abstract

Abnormalities of brain connectivity are consistently observed in individuals with schizophrenias (SZ). Underlying these anomalies, convergent in vivo, post-mortem, and genomic evidence suggest abnormal oligodendrocyte (OL) development and function, including lower *in vivo* myelination in SZ. Using patient derived induced pluripotent stem cells (IPSCs), we previously observed a significant and substantial reduction in the number of OLs produced in cells from the SZ group compared to the healthy control (HC) group. We a lso observed a correlation between white matter (WM) estimated in brain *in vivo* and the number of OLs produced *in vitro*.

We have now characterized potential mediators that may contribute to the SZ-associated deficit in OL production. We ran qRT-PCRs to detect group-specific differences in key myelin pathway proteins. Significant reductions of PAX6 and SOX10 expression were seen in the SZ group. We focused on SOX10 since one of its functions is the commitment of precursor cells to an oligodendrocyte fate. Using an inducible lentiviral system, we expressed SOX10 in patterned neural stem cells (NSCs) and quantified the number of OLs produced. Expression of SOX10 rescued the SZ-associated deficit in OL production, indicating that reduced SOX10 may be a critical mediator of OL dysfunction in SZ. We then ran qRT-PCRs to screen mRNAs for three proteins (SOX9, QK1 and FEZ1) whose expression was directly influenced by SOX10 or directly influenced the expression of SOX10. We saw significant reductions of SOX9 expression and a reduction in QK1 expression in the SZ group. RNAseq analysis confirmed these gene expression changes.

## Introduction

Many genes and environmental factors can contribute to the development of schizophrenias (SZ). To date hundreds of genes have been found that are associated with SZ in large scale Genome Wide Association Studies (GWAS) (reviewed in Hall and Bray, 2022), and it has been predicted that thousands of genes contribute to risk. In rare cases, CNVs (copy number variants) or loss of function mutations can contribute substantially to risk, though they are not fully determinative of illness (Singh et al., 2022). Few factors (variants) contribute more than small individual effects, and determinants or combinations of risk factors underlying single cases can be highly variable across patients. However, some factors, operating in common or convergent cellular pathways, have been consistently observed to be abnormal in SZ. Studies aimed at better understanding these pathways and their controlling elements might guide the development of more effective and better targeted therapeutics.

Many SZ-associated risk variants are in sequences before genes involved in early developmental processes. In part based on this evidence, it has been hypothesized that perturbations of early neural cell development may affect the timing and efficiency of cell maturation, ultimately altering the number and type of cell fate decisions (reviewed by Eyles 2021).

Until recently is has been difficult to study these early developmental processes. Animal models of certain aspects of SZ exist, but neural line development in those animals may not closely mimic the human case. Investigating differences (changes) in post-mortem SZ brain tissue is valuable, but cannot be used to look at questions of early neurodevelopment.

Complementing these approaches, the use of human induced pluripotent stem cells (hIPSCs) models, and the expanding numbers of neural and glial differentiation protocols (Lanjewar and Sloan 2021 and refs therein), provide systems to assess some of the early developmental aspects of SZ. An advantage of using hIPSC models is that the genetic background of the patient is present in the IPS and differentiated cells. These systems can offer an early developmental window into processes that may be dysregulated in SZ, and related diseases, at the first stages of embryonic development.

Among the abnormalities associated with SZ are altered brain developmental pathways that effect connectivity and, consequently, brain signal transduction. Specifically, multiple lines of evidence suggest that white matter abnormalities, associated with abnormal myelination, are typical of SZ. The underlying mechanisms for these deficits are genetic, and ultimately genetically determined developmental factors may directly contribute to altered brain connectivity. This is a set of pathways that we and others have begun to explore.

Using patient derived induced pluripotent stem cells (IPSCs) differentiated to oligodendrocytes, we have previously observed and reported that there is a significant and substantial reduction in the number of OLs produced in the SZ group when compared to OLs produced by the healthy control (HC) group. We also found that there was a correlation between brain imaging estimates of myelination *in vivo* and the number of OLs produced *in vitro*. (McPhie et al 2018). Reduced numbers of OLs were also seen in a separate study of subjects with early onset childhood familial SZ who carried mutations in CSPG4 (de Vrig et al 2018).

Because SZ is in part developmental in nature, looking at early points of the OL pathway might reveal novel intervention targets. To narrow down where in development abnormalities may be occurring, we documented expression levels of key markers (genes) in the oligodendrocyte/myelination pathway. Using the same subject hIPSC lines as in our previous study, we observed significant reduction in expression of PAX6 in NSCs and SOX10 at D45 in the differentiation paradigm. We focused on SOX10 since one of its important functions in human cells is the commitment of precursor cells to an oligodendrocyte fate (Wang et al 2014, Sock and Wegner 2021). We determined whether increasing SOX10 expression in the SZ lines would enhance the numbers of OLs produced by the SZ lines. We found that exogenous expression of human SOX10 in patterned NSCs rescued the deficit in OL production seen in our previous studies. We then looked at expression levels of three mRNAs in the same pathway whose expression levels are influenced by SOX10: SOX9, QK1 and FEZ1 (see Table 1). We found a reduction in SOX9 and QK1 levels but no change in FEZ1 levels. Our study suggests that there are specific early changes occurring in the OL developmental/myelination pathway that may ultimately lead to the reduction in derived OLs produced from cells of our SZ subjects.

**Table 1.** Functions and association with SZ of genes of interest.

| Gene | Relevant functions in OL pathways | Association with SZ | References` |
| --- | --- | --- | --- |
| PAX6 | Fundamental role in regulation of organogenesis and early differentiation of the nervous system. PAX6 controls the balance between NSC self-renewal and neurogenesis | Strober et al 1999 found a moderate association between a high activity variant of PAX6 and paranoid SZ. | Strober et al 1999 |
| OLIG2 | Transcription factor that determines motor neuron and oligodendrocyte differentiation. Expression of OLIG2 in NG2 cells directs it to become an OL and not an astrocyte. | Associations of several SNPs in OLIG2 and SZ. | Komatsu et al 2020 |
| SOX9 | SOX9 functions in Oligodendroglial specification. SOX9 can induce SOX10 expression in OPCs in cooperation with OLIG2 |  | Reiprich et al 2017 |
| SOX10 | Required for OL terminal differentiation and myelination. | SOX10 SNP associated with age of onset in male Han Chinese subjects.<br>SOX10 DNA methylation associated with downregulation of SOX10 and OL dysfunction in SZ in Japanese population. | Yuan et al 2013<br>Iwamoto et al 2005 |
| QKI | Regulator of Oligodendrocyte differentiation and maturation in the brain. SOX10 3' UTR contains QKI binding sites so QKI may directly control SOX10 expression levels. | QKI-7b is downregulated in SZ. May also be functional variants that increase SZ susceptibility. | Aberg et al 2006 |
| FEZ1 | FEZ1 is involved with OL differentiation. It is an adhesion protein. QKI regulates FEZ1 expression. SOX10 can bind and increase FEZ1 expression. | Two SNPs in FEZ1 found in a Japanese SZ cohort. Also, FEZ1 is an interacting partner for DISC1 | Chen et al 2017<br>Yamada et al 2004 |

## Methods

### Research Subjects

Subjects included 6 healthy controls (HC) and 6 patients with schizophrenias, including schizoaffective disorder (together: SZ), recruited from the clinical services at McLean Hospital. Participants were assessed using the Structured Clinical Interview for the DSM-V (SCID)(APA reference) and did not have significant acute or substantial chronic medical illness or conditions. Subjects were studied for brain myelin content in a previous protocol, Du et al 2013. Detailed inclusion and exclusion criteria are given in that reference. There are no previous data on the degree of oligodendroglia production abnormalities in SZ, but sample size was chosen on the basis of the myelin and WM abnormalities previously reported. (Du et al 2013). Our prior expectation was that we would be able to see differences as great as those observed on brain imaging. All protocols were approved by the Partners Healthcare Institutional Review Board, and subjects provided written informed consents before participation in any studies.

### Derivation and expansion of fibroblasts

Fibroblasts were grown from 3mm skin biopsies, cut into small pieces, placed in plate wells with 2ml minimal essential media plus 15% FBS and 1% Penn/Strep under coverslips for 7 days in an incubator at 37ºC, 5% CO^2^. Coverslips were removed when cells were clearly seen migrating out of tissue pieces. Cells were passaged when 80-95% confluent, first to one 100mm dish, then to five 150mm dishes, then frozen and stored. Fibroblast stocks were tested for mycoplasma.

### Conversion to induced pluripotent stem cells

Fibroblasts were sent to either the New York Stem Cell Foundation Research Institute (NYSCF) or Cellular Reprogramming, Inc. (San Diego, CA) for RNA reprogramming. 4SZ and 5HC lines were derived at NYSCF and 2SZ and 1HC lines (converted as part of another study), were from Cellular Reprogramming. NYSCF reprogramming is based on mRNA/miRNA transfection as detailed in Paull et al 2015. Fibroblasts sent to Cellular Reprogramming were similarly converted and the resulting iPS cell colonies were stabilized and expanded. Each fibroblast line was plated to 6-well plates without feeders at three different plating densities and subjected to messenger RNA reprogramming. Colonies were bulk-passaged from the most productive well to establish passage 1 iPS cell cultures on rLaminin-521 (BioLamina) in Nutristem XF media (Biological Industries) and expanded in the same culture system until at least passage 3 before being characterized by DAPI/OCT4/TRA-1-60 immunostaining and frozen down. Cells from both sources were made with the same technology and had similar properties.

### Characterization of iPS cells

After conversion, iPS cell lines from NYSCF were characterized as in Paull etal 2015. Briefly, Nanostring analysis was performed using a 25 gene probe set for markers of pluripotency. Differentiation potential into the three germ layers was assessed using an expression panel of 100 gene probes, after spontaneous differentiation of iPSCs into embryoid bodies. Additionally, cells were karyotyped with the Nanostring nCounter Plex2 Assay Kit. iPSCs reprogrammed at Cellular Reprogramming showed human embryonic stem cell-like morphology and expressed human pluripotent stem cell markers. Characterized iPSCs from both sources were returned to McLean for subsequent studies. iPSCs were mycoplasma tested prior to plating for differentiation.

### qPCRs for developmental markers in the oligodendrocyte pathway

Six SZ and 6 HC lines were differentiated to OLs as in McPhie et al 2018. At D12 2D cultures of NSCs were harvested after Accutase treatment. After the D12 timepoint floating oligospheroids were harvested. Cells or spheroids were pelleted at 300 x g. Media was removed. Pellets were snap frozen on powdered dry ice and stored at -80 C until use.

RNA isolation was done with the RNeasy Plus kit (Qiagen, CA) according to manufacturer’s protocol. cDNA synthesis was performed using Superscript IV VILO kit (ThermoFisher) and 500ng of RNA was used. qRT-PCRs were run to detect expression levels of genes of interest in the oligodendrocyte pathway (see supplemental table 1) using TaqMan gene expression assays and TaqMan fast advanced master mix. Cycling conditions were used based on those recommended for the ViiA7 Real Time PCR system (Applied Biosystems). Control assays for Actin and PPIA were run on each plate for each set of samples.

### Differentiation to Neural Stem Cells for exogenous SOX10 expression

Cells were differentiated to NSCs as in Garcia-Leon, et al 2018 with minor modifications.

Briefly, hIPSCs were plated in E8 (Life Technologies) plus 1uM Y27632 (REPROCELL) at a density of 20,000 cells/cm^2^ (1.3 × 10^5^ cells per well) onto Geltrex (Life Technologies) coated dishes. At the beginning of differentiation medium was changed to N2maxB27 medium (DMEM/F12 supplemented with N2max (1:100), B27 (1:50), glutamax (1:100), NEAA (1:100), beta-mercaptoethanol (1:1000), Pen/Strep (1:100), (all from Life Technologies) and 25 μg/ml insulin (SIGMA). The small molecules SB431542 10uM (REPROCELL), LDN193189 1uM (REPROCELL) and retinoic acid (RA) 100nM (SIGMA) were added from D0-D8. Daily media changes were done until D8. On D9-12 medium was N2maxB27 medium plus 1uM Smoothened agonist (SAG)(Millipore-Sigma) and 100nM RA. On D12 cells were dissociated to single cells with Accutase (Millipore-Sigma) and plated at a density of 1E^6^ cells/well in a 6-well dish, or 4E^5^ cells/well in a 12-well plate. Plates were coated in Poly-L-Ornithine + Laminin (L2020, SIGMA). N2maxB27 medium [RA (100nM), bFGF (20 ng/mL), SAG (1uM)] was the plating medium.

### Lentiviral vectors and virus generation

Lentivirus was produced by Alstem (http://www.alstembio.com/).

### Viral transduction

On the day after plating, cells were transduced with lentivirus at an MOI of 1 based on the number of cells plated. This gives an MOI of 0.5 for each virus used if one cell doubling is assumed from the time of plating. Medium was changed at time of transduction and transduction medium was (N2max+B27 medium [RA (100nM), bFGF (20 ng/mL), SAG (1uM)). Cells were transduced for 16 hr. On the day after transduction doxycycline treatment was begun. Medium was changed -N2maxB27 medium [PDGF (10 ng/mL), HGF (5 ng/mL), IGF-1 (10 ng/mL), NT3 (10 ng/mL), Insulin, Biotin (100 ng/mL), T3 (60 ng/mL), dCAMP (1uM), Doxycycline (2ug/mL)]. Doxycycline treatment was continued for 12 days, media was changed every other day.

### ICC

On D12 of doxycycline treatment O4 positive or SOX10 positive cells were visualized by ICC as in McPhie et al 2018.

### MACS sort of O4 positive cells

Quantification of O4 positive cells was done by magnetic sorting (MACS) following manufacturer’s protocol minor modifications. Briefly cells were dissociated with Accutase (Fisher). Cells were then passed through a 70um filter (Miltenyi). The cells were then pelleted 300g for 5 min. The pellet was resuspended in 90 ul of MACS buffer (PBS + 0.5% BSA). Ten ul of O4 magnetic beads was added and the sample was incubated at 4 degrees for 15min. One ml of MACS buffer was added and samples were added to MS columns (Miltenyi). Negative and positive sort fractions of the sample were collected and cell numbers were quantified.

### CellProfiler analysis of SOX10 positive cells

Images of SOX 10 positive cells from coverslips stained with anti-SOX10 antibody were quantified using CellProfiler image analysis software(https://cellprofiler.org). Briefly SOX10 positive cells and total cells were quantified SOX10 positive cells were expressed as a percentage of total cells.

### RNAseq analysis

Total RNA was sent to Novogene, USA for RNA seq analysis.

### Bioinformatic analysis of transcriptomes

mRNA sequencing data from 24 samples (12 cell lines at day 18 and day 45), after quality control and pre-processing (including demultiplexing, filtering and adapter trimming), were used to detect the association between schizophrenia and gene expression profiles. Raw sequencing reads were mapped to human reference genome (GRCh38) with gapped-alignment strategies. STAR (version 2.7.9a) was used for the alignment and gene-level quantification was performed based on the aligned reads with featureCounts (version 2.0.1). The expression level of a gene in a sample is captured by the number of reads mapped to this gene.

### Differential expression between subjects with schizophrenia (SZ) and healthy controls

Differential expression was detected separately for the two time points using the R package DESeq2(Love et al, 2014). Raw counts were normalized by median of ratio method (Anders and Huber, 2010) to account for artificial variability associated with sequencing depth, gene length and RNA composition. Generalized linear models were applied to the normalized count data to estimate changes in expression level associated with SZ compared to healthy controls for each gene and a Wald test was performed to determine the statistical significance of the estimated change. P-values for different genes within a time point were adjusted with Benjamini-Hochberg (BH) procedure. Bonferroni correction was further applied to BH adjusted *p-values* to account for multiple comparisons in two time points. Our primary hypothesis was that two genes related to myelination pathway (OLIG2, SOX10) would exhibit disease group specific differences in expression. SOX10 and OLIG2 were our main targets in the analysis. Exploratory analysis was also performed on an extended list of genes (see supplementary materials). The significance level for all the tests is set at 0.05.

### Gene-set enrichment analysis

In addition to identify individual genes that are differentially expressed in patients with SZ, gene-sets of interest were also examined with Gene-set enrichment analysis (GSEA) (Subramanian et al 2005; Mootha et al 2003). Three self-defined gene-sets based on Sock and Wegner 2021 were considered: 1) genes related to myelination pathway, including OLIG2, NKX2-2, SOX9, SOX10, QK1, FEZ1 and TCF7L2; 2) OLIG2 gene-set, including SOX9, SOX10, MYRF, SOX5, SOX6, ZEB2, SMARCA4, ID1, TCF4, ZNF24, CHD8 and ZNF488; 3) SOX10 gene-set, including NKX2-2, QK1, FEZ1, TCF7L2, MYRF, SOX6, ZNF24, HES5, NFATC2, PDGFRA, CSPG4 and HIF1A. Data from the two time points were analyzed separately, BH adjusted p-values were reported for the statistical significance of the observed enrichment/depletion. The significance level for all the tests is set at 0.05.

### Association of single-nucleotide variants (SNVs) in exome to SZ

Whole exome sequencing (WES) was performed for 11 out of 12 subjects from whom the cell lines are derived. (One control subject’s DNA was omitted from the much larger set that was exome sequenced). SNVs were detected using the Broad/GATK pipeline for calling (GATK Team, 2022). All sequencing data, including those generated externally and data generated in earlier analyses, are centralized as BAM or FASTQ files, and processed uniformly using Picard sequence processing pipeline, before being mapped onto the human genome reference build 37 (grch37) using BWA (Li H. and Durbin R., 2010). A combination of GATK v3.4 and v3.6 (Van der Auwera and O’Connor, 2020) was then used for calling and VerifyBamID (Jun et al, 2012) version 1.0.0 for sequence analysis. Regular microarray data was available for the remaining subject and was combined with the WES data for the downstream analysis. SNVs that fall within the regions of the genes PAX6, SOX1, NKX2-2, OLIG2, SOX10, SOX9, QKI and FEZ1 were considered. PLINK2 was used to perform the statistical tests. Due to the small sample size in our dataset, only sex was included as the covariate in the regression model and population substructure was ignored in the analysis. Since this analysis was mainly for exploratory purpose and the sample size is limited, the p-values were not adjusted for multiple comparisons and the significance level is set at 0.25.

## Results

### qRT-PCR analysis of OL developmental markers in our oligodendrocyte differentiation protocol

To define the point(s) of the oligodendrocyte developmental pathway where dysregulation is occurring in our SZ subjects, we performed a series of qRT-PCRs of known regulators of oligodendrocyte development at time points when these genes are known to be expressed in this differentiation protocol (Douvaras et al 2014). PAX6 and SOX1 were tested at D8, and OLIG2 and NKX 2.2 at D12. SOX10 was tested at D45 and D50.

We saw a significant difference in PAX6 expression at D8 of the differentiation. SOX 1 showed no difference in expression at that timepoint. Also, no significant differences in expression of NKX 2.2 was seen at D12, indicating that at this early step of development not all components of the pathway were dysregulated. OLIG2 expression at D12 by qPCR showed increased expression in the SZ group, but this increase did not reach statistical significance at this early timepoint. SOX 10 expression was examined at D45 and D50, based on gene expression data in Douvaras and Fosatti 2015, and because at these timepoints we see SOX10 positive cells emerging from the oligospheres. A significant reduction in SOX10 expression was seen in the SZ group, in comparison to healthy controls, at both time points (Fig 1). qRT-PCR results were repeated on cells from two independent differentiations.

**Fig 1.**
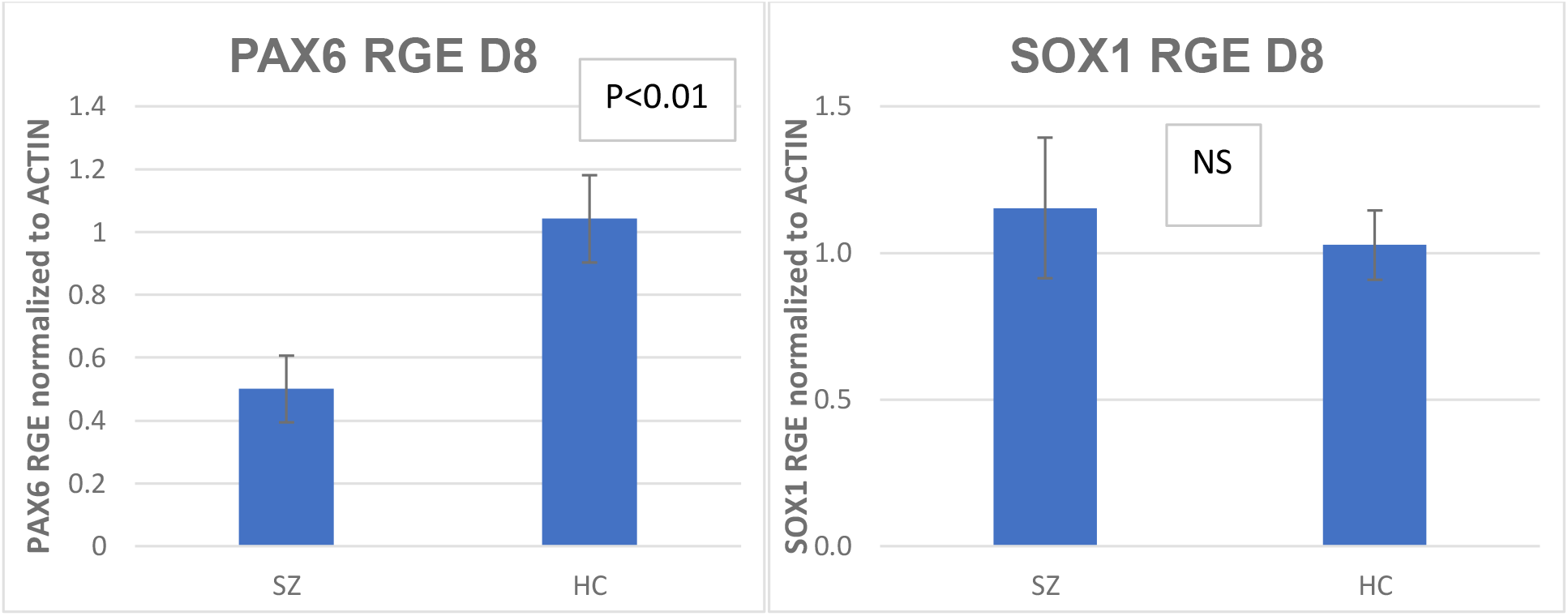
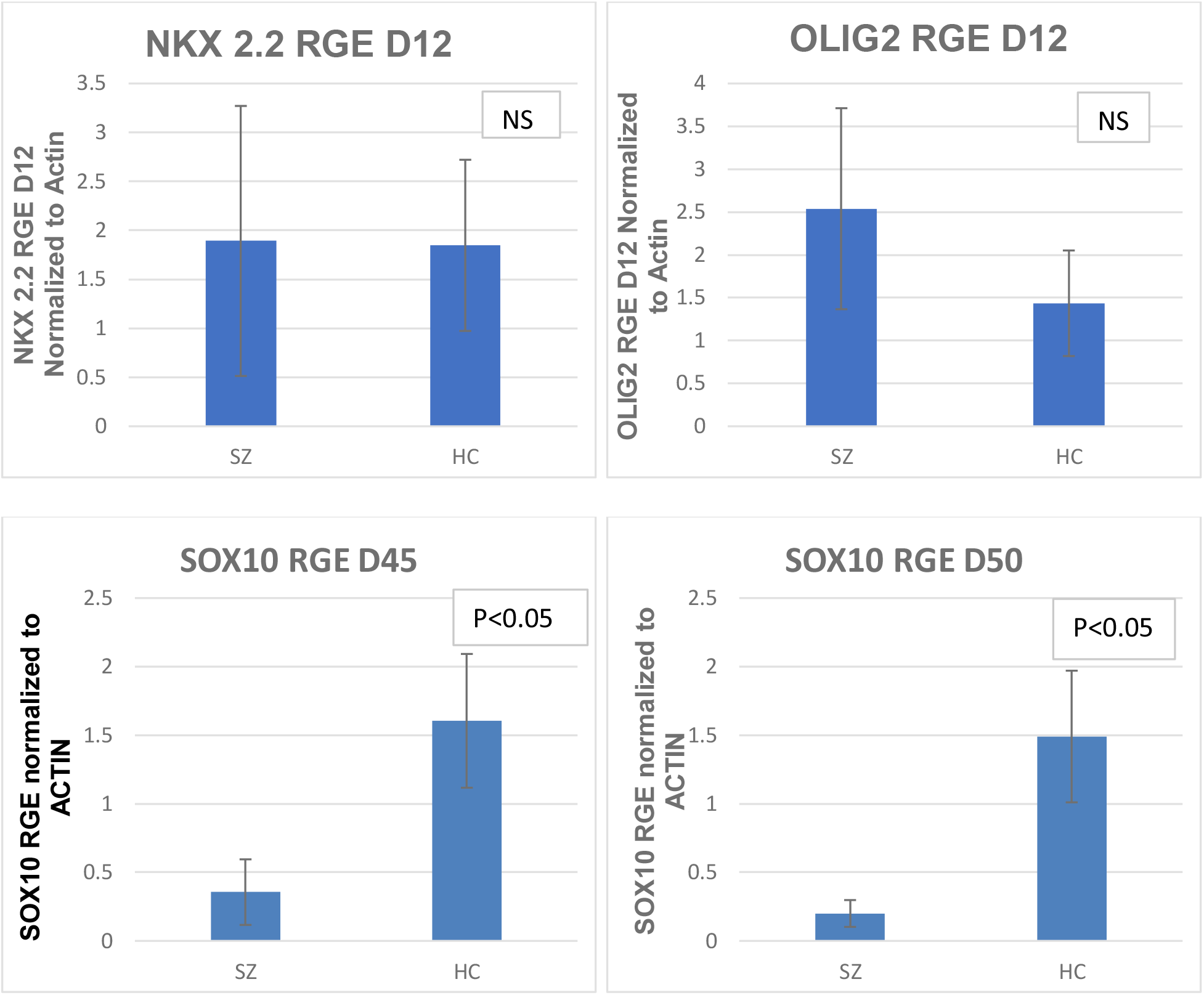
Differential gene expression of key genes in the myelination pathway in derived cells from SZ and control subjects. qRT-PCRs show disease specific differences in PAX6 (p<0.01), OLIG2 and SOX10 (p<0.05) expression. SOX1 and NKX2.2 showed no disease specific expression differences.

### Exogenous expression of SOX10 with lentivirus rescues the OL developmental deficit seen in the SZ subject lines

Since SOX10 is critical for differentiation and maturation of OLs, we determined whether increasing the expression level of SOX10, using an exogenous source of SOX10 delivered through an inducible lentiviral vector system, would increase the number of OLs produced in the SZ subject lines, allowing them to overcome the disease specific deficit we previously observed (McPhie et al 2018). Lentiviral vectors were added to D13 patterned NSCs. One day after viral transduction doxycycline was added to the medium to induce expression of SOX10 in the NSCs. Twelve days after induction with doxycycline, the cultures were harvested and O4 positive OPCs and OLs were assessed by MACS sorting. We found no significant differences in the percentage of O4 positive cells in the SZ (n=6) vs the HC group (n=6). This indicated that the defect in OL differentiation pathway was, at least substantially, at the point of SOX10 expression or upstream of this point in the pathway. (Fig 2) NSCs infected with the lentiviruses and not induced by doxycycline showed 1% or fewer of O4 positive cells. Additionally, we examined the percentage of SOX10 positive cells by ICC in each line to control for line-to-line variability in expression. We found a mean of 42.5% for the SZ group and a mean of 42.6% for the HC group with a range of 33 to 55% for HC origin lines and 32-48% for SZ origin lines. That is, the effect associated with SOX10 expression was observed in all of the SZ origin lines. An example image of the SOX10 expression in the transduced cells from SZ and HC groups is shown in Fig 2C. The SOX10 rescue experiment was repeated in 2 independent differentiations of the same subject lines.

**Fig 2.**
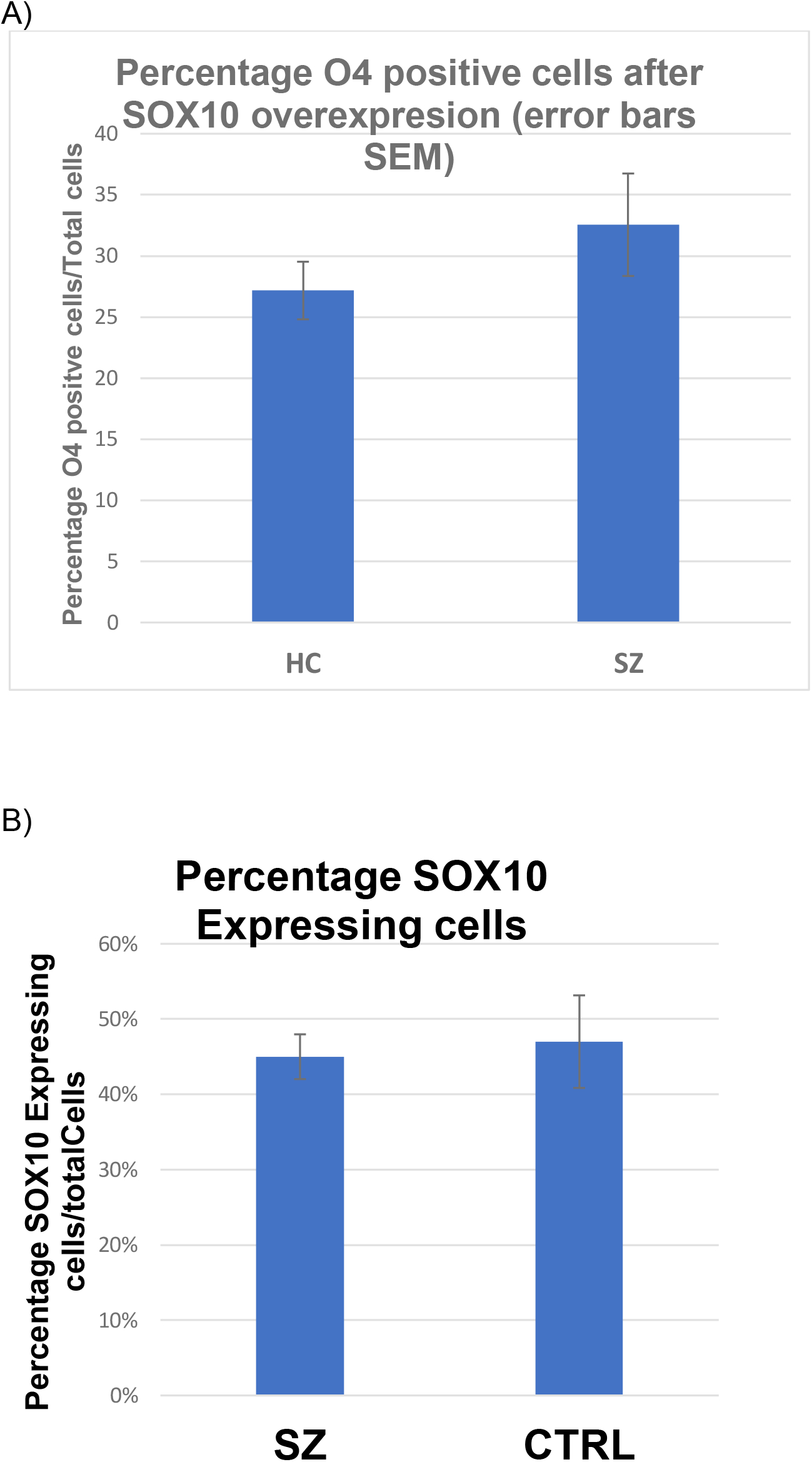
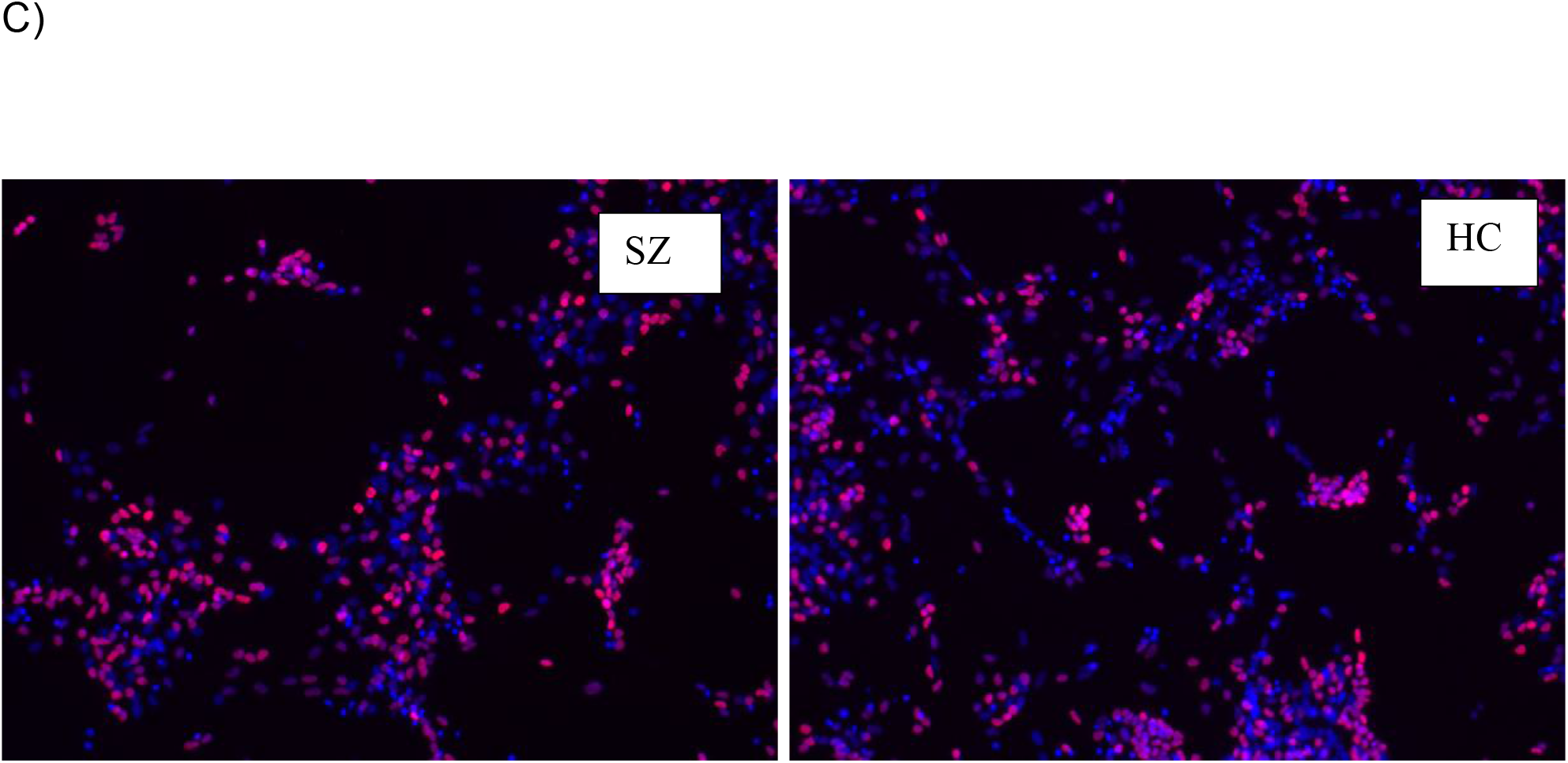
Exogenous expression of SOX10 in patterned neural stem cells recues the SZ deficit in OL production. A) MACS sort data for O4 positive cells. B) ICC quantification of SOX10 expression in transduced cells. C) ICC of patterned NSCs 12 days after transduction to show exogenous SOX10 expression levels SOX10 (red) and Hoechst (blue)

### Analysis of the expression levels of SOX9, QKI and FEZ1 by qRT-PCR

Using qRT-PCR, we determined the expression of key myelin pathway markers (QKI, FEZ1 and SOX9) in spheres at D50 of our standard differentiation protocol. These genes have been shown to either influence the expression of SOX10 or whose expression is influenced by SOX10. SOX9 has been shown to induce SOX10 expression in OPCs in cooperation with OLIG2 (Reirprich et al 2017). SOX10 contains QKI binding sites in its 3’ UTR so QKI may directly control SOX10 levels (Aberg et al. 2006). For FEZ1, SOX10 has been shown to increase FEZ1 expression levels (Yamada et al 2004). Table 2 summarizes these relationships and also lists the association of these genes to SZ.

We found that SOX9 showed a significant reduction in expression in the SZ group (p<0.05) vs controls and that QK1 showed a strong trend of reduced expression in the SZ subjects vs controls (p=0.05). No group specific differences were seen in FEZ1 expression levels. (Fig 3)

**Fig 3.**
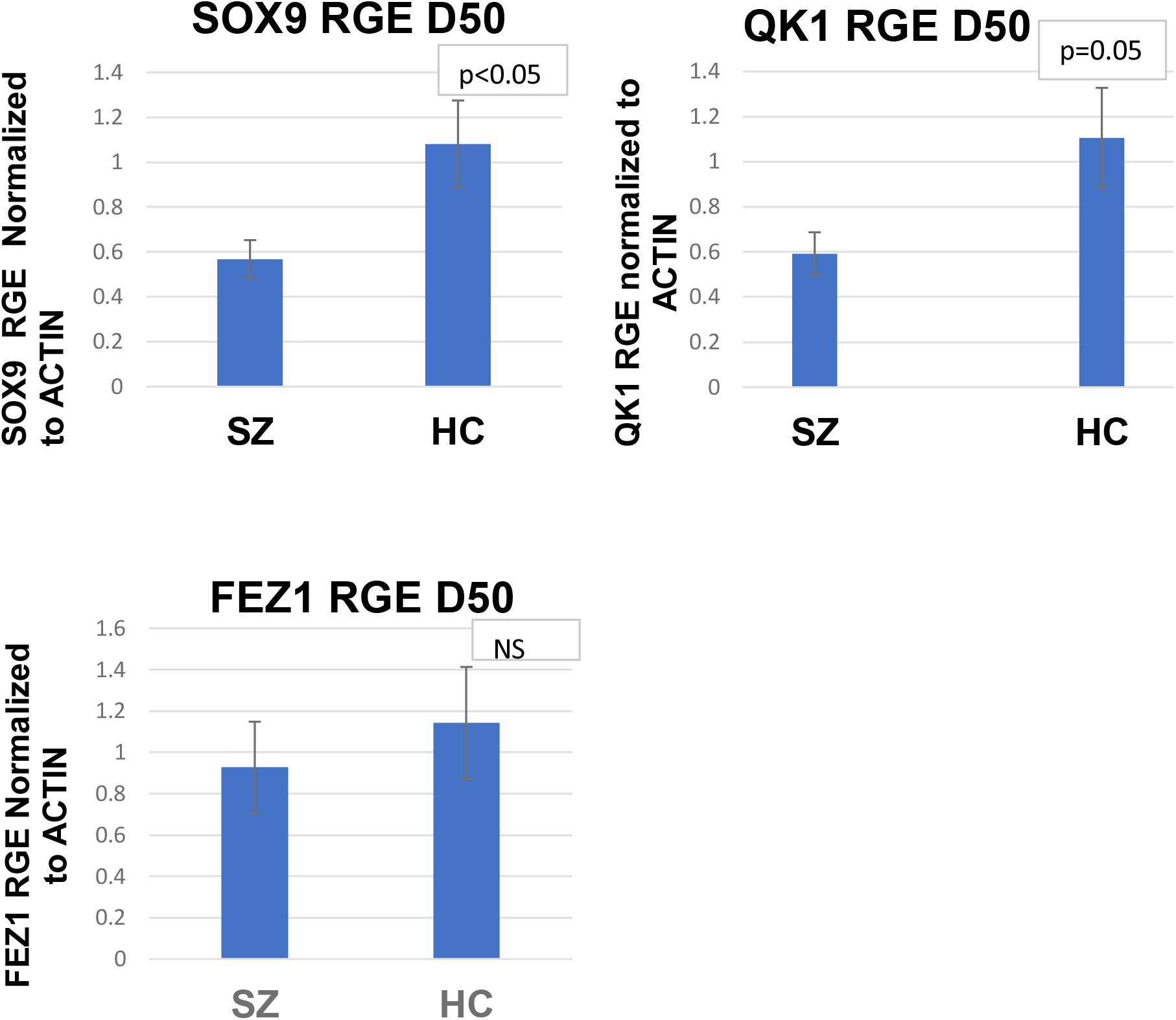
Differential Gene Expression of Three Genes Known to be influence or be influenced by SOX 10 Expression in cells from the SZ and HC Group. Disease specific differences in expression were seen for SOX9 (p<0.05) and QKI (p=0.05) using cells from D50 of our standard differentiation. FEZ1 did not show disease specific differences.

### RNA seq analysis

#### Differential expression of genes related to myelination pathway in SZ

The results of the differential expression analysis for SOX10 and OLIG2 at day 18 and day 45 are shown in Figure 4 A and B. The results at D18 suggest that SOX10 is significantly up-regulated in SZ compared to healthy controls and OLIG2 is also up-regulated in SZ, although the association is not significant. At day 45, OLIG2 is significantly upregulated in SZ compared to healthy controls and SOX10 is significantly downregulated. A list of additional genes related to myelination were also tested in an exploratory analysis. Three additional genes – SOX9, CSPG4 and ZFP488 – are significantly upregulated in patients with SZ at day 18 while none of the genes in the list have significantly different expression profiles in SZ at day 45. (Supplementary table 2)

**Fig 4.**
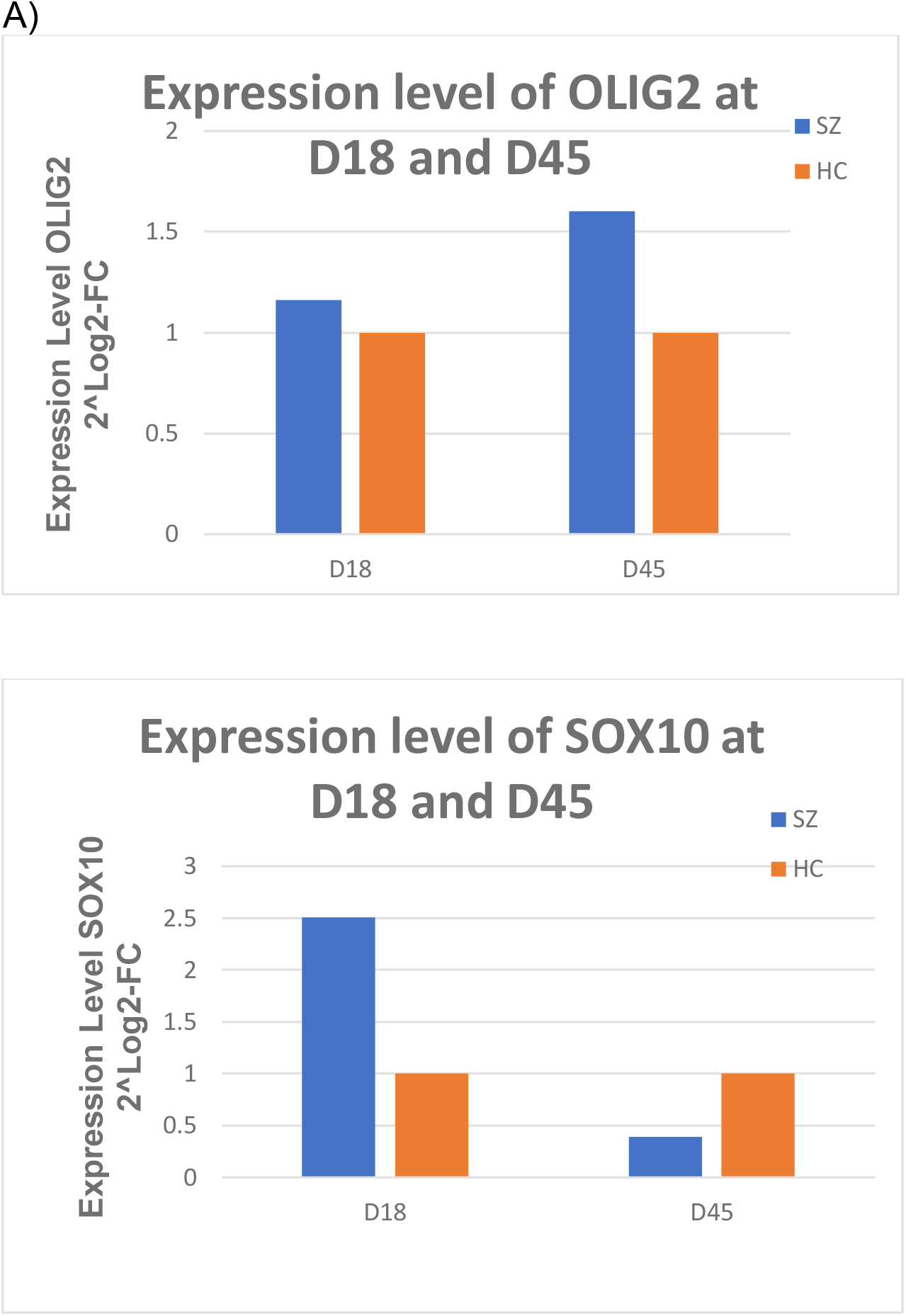
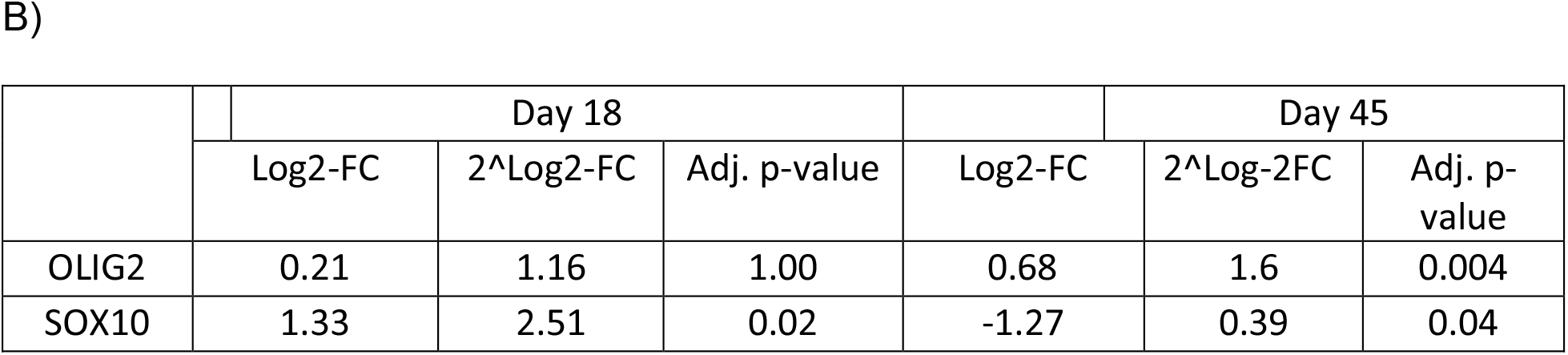
RNAseq Analysis of OLIG2 and SOX10 differential expression by time for the SZ and HC groups. At D18 the data suggest that SOX10 is significantly up-regulated in SZ compared to healthy controls whereas OLIG2 is also up-regulated in SZ b. At day 45, OLIG2 is significantly upregulated in SZ compared to healthy controls and SOX10 is significantly downregulated.

#### GSEA analysis of self-defined gene-sets

GSEA results indicate that all three gene-sets are up-regulated in SZ compared to healthy controls at day 18. At day 45, gene-set related to myelination pathway becomes down-regulated in SZ and the directionality of enrichment remains the same for the other two gene-sets. However, none of the enrichment results are statistically significant. See Supplementary Table 3 and Supplementary Figure 1 for details.

#### Association between SNVs near target genes and SZ

When focusing on SNVs in the exomes near the target genes, weakly significant results are observed for three SNPs: rs2071754 (p-value = 0.13), rs1352461141 (p-value = 0.19) and rs1791018024 (p-value = 0.21). The first two SNPs are in gene PAX6, one is an intronic variant (rs2071754) and the other a synonymous variant (rs1352461141).

The third SNP is in gene QKI and is an intronic variants. Both of these genes have been associated with SZ and variants in area of the exome near the target may affect their expression level.

## Discussion

The results reveal specific abnormalities in the early stages of OL development in reprogrammed cells from subjects with SZ. Specifically, there are differences in expression of PAX6, OLIG2, and SOX10 in the OL developmental pathway in SZ derived cells. However, the expression of other genes in the OL developmental pathway is unchanged, indicating a more complicated story than simply that the entire OL differentiation pathway is down-regulated in SZ. When we focused on SOX10 expression levels and manipulated these in both SZ and HC groups, using an inducible lentiviral expression system, we were able to rescue the deficit in OLs observed in our SZ subject lines. When then examined three key myelin pathway markers, for gene products interacting with SOX10 (SOX9, QK1 and FEZ1), at day 50. We found decreased expression of SOX9 and QK1 but no difference in FEZ1 levels, providing further evidence that only some specific markers are reduced in our SZ subjects.

PAX6 is a high-level regulatory gene that plays a role in neurogenesis and precursor proliferation (Manual et al 2015). The difference in PAX6 expression between the two diagnostic groups occurs at a point very early on in the differentiation process, when the cells have not yet specified a commitment to a neuronal or glial fate pathway. Because PAX6 expression is reduced but SOX1, another marker of NSCs, expression is unchanged the reduction in PAX6 levels is most likely not just due to a decrease in total NSCs produced by the SZ subject’s cells at this point in development. There have been other studies that implicate a role for PAX6 in SZ. Stober et al 1999 found a moderate association of a high activity variant of PAX6 linked to paranoid SZ. These authors hypothesize that the PAX6 polymorphism may me a modifier of individual SZ risk. In contrast Brennand et al 2015 and Liu et al 2019 did not find PAX6 expression differences in NPCs from childhood onset SZ subjects. Both used different differentiation strategies and protocols, so that may account for the difference in results. SZ is associated with abnormalities in more than one neuronal and more than one glial cell type. Thus, reduced expression of PAX6 may underlie reductions of any of these mature neurons and glia.

We present detailed evidence that this cell model might be used to clarify one particular pathophysiologic mechanism, abnormal myelination, contributing to the risk and development of SZ. In time, it might provide a platform to test methods or agents for amelioration or repair.

In humans SOX10 protein controls the commitment and induction of glial precursor cells to an oligodendrocyte fate (Wang et al 2014 Sock and Wegner 2021). Later in development it serves a role in terminal differentiation of oligodendrocytes and, through the mature functions of those cells, myelination. Many studies have reported down regulation of OL genes in SZ (Valdes-Tovar et al, 2021; Haroutunian et al, 2014; Roussos and Haroutunian 2014). Two studies have found partial associations of SOX10 and SZ. One study by Iwamoto et al 2005 found a correlation between a decrease in SOX10 expression and the degree of methylation of the CpG island of the SOX10 gene in brains of SZ subjects. The increased methylation was specific to SOX10 and not found in other OL specific regulators such as OLIG2 or MOBP (Iwamoto et al 2005). Yuan et al 2013 reported an association of the SOX10 rs139883 polymorphism and age of onset of SZ in a Han Chinese population (Yuan et al 2013).

To our knowledge no other studies have examined the outcome of modification of SOX10 levels in human derived cells from subjects with SZ. A study using a Huntington’s disease (HD) chimeric mouse model (Osipovitch et al 2019) examined glial pathology in HD and observed that forced expression of both SOX10 and MYRF restored the myelination defect seen in this model. The effects of SOX10 expression, alone, was not reported. However, in combination with the results reported here, the findings suggest that glial growth enhancement by targeting developmental factors may be a viable therapeutic avenue.

Additional experiments are needed to define the role of SOX10 and its interactions with other developmental factors. OLIG2 is known to induce SOX10 through a distal enhancer. (Kuspert et al., 2011; Sock and Wegner 2021) As more information becomes available, it will be interesting to learn if OLIG2 builds up over time in SZ lines because OLIG2 or other interacting factors do not effectively induce SOX10 levels in SZ. Nonetheless, the findings help define key potential targets for intervention in a very complex developmental pathway. Limitations of the study include a modest number of subject lines in each group and the use of a lentiviral system to exogenously control gene expression to speed up the OL developmental pathway in human derived NSCs.

We present evidence that increasing SOX10 expression at an early point of the pathway may overcome the deficit of decreased OL production. Correcting this deficit may ultimately help in normalizing white matter levels. However, such intervention might only be effective during brain development, in utero. Also, increasing SOX10 expression levels might alter other downstream processes in a way that would not ultimately benefit the patient.

Of note, we are not suggesting that reduced OL development and consequent white matter deficits explain more than a portion of the pathophysiology underlying SZ. The OL development/myelination pathway does however seem to be a key determinant of illness. And not all illness related abnormalities might need to be addressed to ameliorate symptoms. With more study, oligodendrocyte number may be a tractable target for therapeutic development *in vivo*.

## References

Aberg K, Saetre P, Jareborg N, Jazin E. Human QKI, a potential regulator of mRNA expression of human oligodendrocyte-related genes involved in schizophrenia. Proc Natl Acad Sci U S A 2006; 103: 7482–7487. doi: 10.1073/pnas.0601213103.Epub 2006 Apr 25.

Anders S, Huber W. Differential expression analysis for sequence count data. Genome Biol. 2010; 11: 106. http://dx.doi.org/10.1186/gb-2010-11-10-r106

APA. Diagnostic and statistical manual of mental disorders (5th ed.) American Psychiatric Publishing: Arlington, VA, 2013.

Brennand K, Savas JN, Kim Y, Tran N, Simone A, Hashimoto-Torii K, et al. Phenotypic differences in hiPSC NPCs derived from patients with schizophrenia. Mol. Psychiatry. 2015; 20: 361–8. doi: 10.1038/mp.2014.22. Epub 2014 Apr 1. PMID: 24686136; PMCID: PMC4182344.

Chen X, Ku L, Liu G, Xu Z, Wen Z, Zhao X, Wang F, Xiao L, Feng Y. Novel schizophrenia risk factor pathways regulate FEZ1 to advance oligodendroglia development. Transl. Psychiatry 2017; 7: 1293. DOI 10.1038/s41398-017-0028-z

de Vrij, FM, Bouwkamp CG, Gunhanlar N, Shpak G, Lendemeijer B, Baghdadi M et al. Candidate CSPG4 mutations and induced pluripotent stem cell modeling implicate oligodendrocyte progenitor cell dysfunction in familial schizophrenia. Mol. Psychiatry 2019; 24: 757–771. https://doi.org/10.1038/s41380-017-0004-2.

Douvaras P, Fossati V. Generation and isolation of oligodendrocyte pro-genitor cells from human pluripotent stem cells. Nat. Protoc. 2015; 10: 1143–1154.

Douvaras P, Wang J, Zimmer M, Hanchuk S, O’Bara MA, Sadiq S, et al. Efficient generation of myelinating oligodendrocytes from primary progressive multiple sclerosis patients by induced pluripotent stem cells. Stem Cell Rep. 2014; 3: 250–259. http://dx.doi.org/10.1016/j.stemcr.2014.06.012

Du F, Cooper AJ, Thida T, Shinn AK, Cohen BM, Ongur D. Myelin and axon abnormalities in schizophrenia measured with magnetic resonance imaging techniques. Biol Psychiatry 2013; 74: 451–457.

Eyles, DW. How do established developmental risk-factors for schizophrenia change the way the brain develops? Transl Psychiatry 2021; 11: 158. https://doi.org/10.1038/s41398-021-01273-2

Garcia-Leon JA, Kumar M, Boon R, Chau D, One J, Wolfs E, et al. SOX10 single transcription factor-based fast and efficient generation of oligodendrocytes from human pluripotent stem cells. Stem Cell Rep. 2018; 10: 1–18 https://doi.org/10.1016/j.stemcr.2017.12.014

GATK Team. Germline short variant discovery. 2022. Available: https://gatk.broadinstitute.org/hc/en-us/articles/360035535932-Germline-short-variant-discovery-SNPs-Indels-

Hall J, Bray NJ. Schizophrenia genomics: convergence on synaptic development, adult synaptic plasticity or both. Biol Psychiatry 2022; 91: 709–719.

Haroutunian V, Katsel P, Roussos P, Davis KL, Altshuler LL, Bartzokis. Myelination, oligodendrocytes, and serious mental illness. Glia 2014; 62:1856–1877.

Iwamoto K, Bundo M, Yamada K, Takao H, Iwayama-Shigeno Y, Yoshikawa T, Kato T. DNA methylation status of SOX10 correlates with its downregulation and oligodendrocyte dysfunction in schizophrenia. J Neurosci. 2005; 25: 5376–81. doi: 10.1523/JNEUROSCI.0766-05.2005. PMID: 15930386

Jun G, Flickinger M, Hetrick KN, Romm JM, Doheny KF, Abecasis G, et al. Detecting and Estimating Contamination of Human DNA Samples in Sequencing and Array-Based Genotype Data. Am. J. of Hum. Genet. 2012; 91: 839–848. doi:10.1016/j.ajhg.2012.09.004

Komatsu H, Takeuchi H, Kikuchi Y, Ono C, Yu Z, Iizuka K, et al. Ethnicity-dependent effects of schizophrenia risk variants of the OLIG2 gene on OLIG2 transcription and white matter integrity. Schizophr. Bull. 2020; 46:1619–1628. Published online 2020 Apr 14. doi: 10.1093/schbul/sbaa049 PMCID: PMC7846078

Küspert M, Hammer A, Bösl MR, & Wegner M. Olig2 regulates Sox10 expression in oligodendrocyte precursors through an evolutionary conserved distal enhancer. Nucleic Acids Res. 2011; 39: 1280–1293.

Lanjewar SN and Sloan SA. 2021. Growing glia: cultivating human stem cell models of gliogenesis in health and disease. Front. Cell Dev. Biol. 2021; 9: 649538. Doi: 10.3389/fcell.2021.649538

Li H, Durbin R. Fast and accurate long-read alignment with Burrows-Wheeler Transform. Bioinformatics 2010, Epub. [PMID: 20080505]

Liu Z, Osipovitch M, Benraiss A, Huynh NPT, Foti R, Bates J, et al. Dysregulated glial differentiation in schizophrenia may be relieved by suppression of SMAD4-and REST-dependent signaling. Cell Rep. 2019; 27: 3832–3843.e6. doi: 10.1016/j.celrep.2019.05.088. PMID: 31242417; PMCID: PMC6700735.

Love MI, Huber W, Anders S. Moderated estimation of fold change and dispersion for RNA-seq data with DESeq2. Genome Biology 2014

Manuel MN, Mi D, Mason JO, Price DJ. Regulation of cerebral cortical neurogenesis by the Pax6 transcription factor. Front. Cell. Neurosci. 9: 70. doi: 10.3389/fncel.2015.00070

McPhie DL, Nehme R, Ravichandran C, Babb SM, Ghosh SD, Staskus A, et al. Oligodendrocyte differentiation of induced pluripotent stem cells derived from subjects with schizophrenias implicate abnormalities in development. Transl. Psychiatry 2018; 8: 230. doi: 10.1038/s41398-018-0284-6

Mootha V, Lindgren C, Eriksson KF, Subramanian A, Sihag S, Lehar J et al. PGC-1α-responsive genes involved in oxidative phosphorylation are coordinately downregulated in human diabetes. Nat. Genet. 2003; 34:267–273. https://doi.org/10.1038/ng1180

Osipovitch M, Asenjo Martinez A, Mariani JN, Cornwell A, Dhaliwal S, Zou L, et al. Human ESC-Derived chimeric mouse models of Huntington’s Disease reveal cell-intrinsic defects in glial progenitor cell differentiation. Cell Stem Cell. 2019; 24: 107–122.e7. doi: 10.1016/j.stem.2018.11.010. Epub 2018 Dec 13. PMID: 30554964; PMCID: PMC6700734.

Paull D, Sevilla A, Zhou H, Hahn AK, Kim H, Napolitano C et al. Automated, high-throughput derivation, characterization and differentiation of induced pluripotent stem cells. Nat. Methods 2015; 12: 885–892.

Reiprich S, Cantone M, Weider M, Baroti T, Wittstatt J, Schmitt C, Kuspert M, Vera J, Wegner M. Transcription factor Sox10 regulates oligodendroglial Sox9 levels via microRNAs. Glia 2017; 65: 1089–1102.

Roussos P. and Haroutunian V. Schizophrenia: susceptibility genes and oligodendroglial and myelin related abnormalities. Front. Cell. Neurosci. 2014; 8: 5.

Schizophrenia Working Group of the Psychiatric Genomics Consortium. Bio-logical insights from 108 schizophrenia-associated genetic loci. Nature 2014; 511: 421–427.

Singh T, Poterba T, Curtis D, Akil H, Al Eissa M, Barchas JD et al. Rare coding variants in ten genes confer substantial risk for schizophrenia. Nature 2022; 604: 509–516. https://doi.org/10.1038/s41586-022-04556-w

Sock E, Wegner M. Using the lineage determinants Olig2 and Sox10 to explore transcriptional regulation of oligodendrocyte development. Dev. Neurobiol. 2021; 81: 892–901.

Stober G, Syagailo YV, Okladnova O, Jungkunz G, Knapp M, Beckmann H, Lesch KP. Functional PAX-6 Gene Linked Polymorphic region: Potential Association with Paranoid Schizophrenia. Biol. Psychiatry 1999; 45: 1585–1591.

Subramanian A, Tamayo P, Mootha VK, Mukherjee S, Ebert BL, Gillette MA, et al. Gene set enrichment analysis: a knowledge-based approach for interpreting genome-wide expression profiles. Proc. Natl. Acad. Sci. U S A. 2005; 102: 15545–15550. doi: 10.1073/pnas.0506580102. Epub 2005 Sep 30. PMID: 16199517; PMCID: PMC1239896.

Valdés-Tovar M, Rodríguez-Ramírez AM, Rodríguez-Cárdenas L, Sotelo-Ramírez CE, Camarena B, Sanabrais-Jiménez MA, et al. Insights into myelin dysfunction in schizophrenia and bipolar disorder. World J. Psychiatry 2021; 12: 264–285.

Van der Auwera GA, O’Connor BD. Genomics in the Cloud: Using Docker, GATK, and WDL in Terra (1st Edition). O’Reilly Media 2020.

Wang J, Pol SU, Haberman AK, Wang C, O’Bara MA, Sim FJ. Transcription factor induction of human oligodendrocyte progenitor fate and differentiation. Proc Natl Acad Sci U S A. 2014; 111: E2885–94. doi: 10.1073/pnas.1408295111. Epub 2014 Jun 30.

Yamada K, Nakamura K, Minabe Y, Iwayama-Shigeno Y, Takao H, Toyota T, et al. Association analysis of FEZ1 variants with schizophrenia in Japanese cohorts. Biol. Psychiatry 2004; 56: 683–690.

Yuan A, Li W, Yu T, Zhang C, Wang D, Liu D, et al. SOX10 rs139883 polymorphism is associated with the age of onset in schizophrenia. J. Mol. Neurosci. 2013; 50: 333–338. DOI 10.1007/s12031-013-9982-y

