## Supplemental figure 1, Supplemental Tables 1, 2 and 3 for "SOX10 Expression Levels may be a Critical Mediator of White Matter Dysfunction in Schizophrenia"

**Supplementary Figure 1 Enrichment score plots from GSEA.**

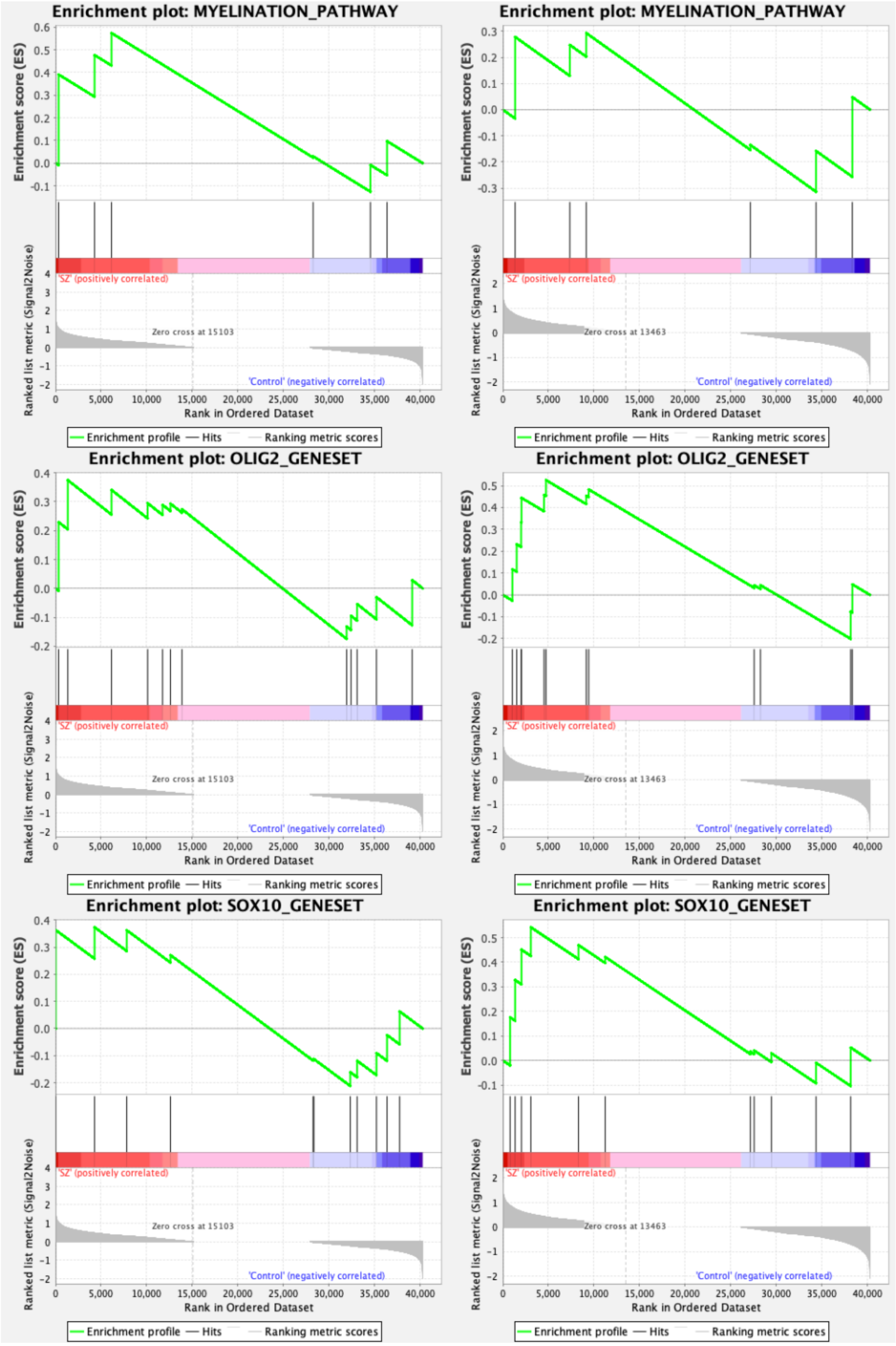

**Supplementary Table 1**

| **TAQman Assays(qPCR primers) used:** |
|---|
| PAX6 Hs0108811(4)4_m1 |
| SOX1 Hs01057642_s1 |
| OLIG2 Hs00377820_m1 |
| NKX2.2 Hs00159616_m1 |
| SOX10 Hs00366918_m1 |
| QK1 Hs 00916678_m1 |
| SOX9 Hs00165814_m1 |
| FEZ1 Hs00363763_m1 |
| ACTIN Hs01060665_g1 |
| PPIA Hs04194521_s1 |

**Supplementary Table 2: Differential expression results from DESeq2 for additional genes related to myelination pathway. FC: fold-change of schizophrenia vs. healthy controls**.

| | Day 18 | | Day 45 | |
|---|---|---|---|---|
| | Log2-FC | Adj. p-value | Log2-FC | Adj. p-value |
| SOX9 | 1.43 | $4.70 \times 10^{-5}$ | 0.69 | 0.10 |
| CSPG4 | 1.44 | $5.05 \times 10^{-4}$ | -0.43 | 0.11 |
| ZFP488/ZNF488 | 5.02 | $3.32 \times 10^{-3}$ | 0.67 | 0.12 |
| ZFP24/ZNF24 | -0.13 | 0.15 | 2.50 | 0.12 |
| TCF7L2 | 0.66 | 0.36 | 0.32 | 0.12 |
| SIP1/ZEB2 | -0.65 | 0.36 | 0.30 | 0.28 |

| | | | | |
|---|---|---|---|---|
| HES5 | -0.39 | 0.64 | 0.63 | 0.28 |
| BRG1/SMARCA4 | 0.10 | 0.72 | 0.53 | 0.28 |
| CHD8 | -0.06 | 0.83 | -0.89 | 0.28 |
| NFATC2 | 0.43 | 0.87 | -0.23 | 0.33 |
| TCF4 | 0.16 | 0.88 | -0.24 | 0.45 |
| SOX5 | 0.21 | 0.88 | 1.46 | 0.86 |
| NKX2.2 | -1.02 | 0.88 | 0.43 | 0.88 |
| QK1 | -0.04 | 1.00 | -0.72 | 1.00 |
| ID1 | -0.38 | 1.00 | -0.19 | 1.00 |
| MYRF | -0.13 | 1.00 | 0.35 | 1.00 |
| SOX6 | 0.24 | 1.00 | -0.05 | 1.00 |
| HIF1A | -0.05 | 1.00 | -0.31 | 1.00 |
| FEZ1 | 0.06 | 1.00 | -0.05 | 1.00 |
| PDGFRA | 0.02 | 1.00 | 0.17 | 1.00 |

**Supplementary Table 3: GSEA results for the three self-defined gene-sets at day 18 and day 45. ES: enrichment score in SZ vs. healthy controls.**

| Gene-set | Day 18 | | Day 45 | |
|---|---|---|---|---|
| | ES | Adj. p-value | ES | Adj. p-value |
| MYELINATION_PATHWAY | 0.57 | 0.75 | -0.31 | 0.94 |
| OLIG2_GENESET | 0.37 | 0.68 | 0.54 | 0.27 |
| SOX10_GENESET | 0.37 | 0.60 | 0.53 | 0.36 |
